# Racial Disparities in the Genetic Landscape of Acute Myeloid Leukaemia from The Cancer Genome Atlas: Insights from a Bioinformatics Analysis

**DOI:** 10.1101/2023.11.06.565754

**Authors:** Panji Nkhoma, Kevin Dzobo, Doris Kafita, Geoffrey Kwenda, Sody Munsaka, Sinkala Musalula

## Abstract

Acute myeloid leukaemia (AML) is a heterogeneous disease with complex pathogenesis that affects hematopoietic stem cells. Ethnic and racial disparities have been reported to affect treatment and survival outcomes in AML patients. Here, we analysed clinical and transcriptomic data from The Cancer Genome Atlas (TCGA) to investigate potential differences in the genetic landscape of AML between African and European individuals. We found several differentially expressed mRNA transcripts between the AML of Africans and Europeans. Notably, AML in African patients exhibited enrichment for several pathways, including signalling by G-protein-coupled receptors, oncostatin M, and codeine and morphine metabolism. In contrast, AML in European patients showed enrichment for pathways related to the glial cell-line derived neurotrophic factor/rearranged during transfection signalling axis, gamma-aminobutyric acid receptor activation, and ligand-gated ion transport channels. Additionally, kinase enrichment analysis identified shared and distinct kinases in AML among Africans and Europeans: Africans showed an enrichment of cyclin-dependent kinases, while Europeans exhibited an enrichment of ULK2, CSNK2B, and CAMK1. Our study highlights the potential importance of considering race when evaluating the genetic landscape of AML, which may improve treatment strategies for this disease.

## Introduction

Acute Myeloid Leukaemia (AML) is a type of blood cancer affecting haemopoietic stem cells, with a complex pathogenesis due to gene mutations, chromosomal abnormalities, and epigenetic changes leading to an increase in non-functional myeloblasts [1, 2]. The incidence of AML has increased globally, with an age-standardized incidence rate rising from 1.35/100,000 in 1990 to 1.54/100,000 in 2017, showing an average percentage change of 0.56 [3]. The application of next-generation sequencing (NGS) technologies has significantly expanded our knowledge of the molecular diversity of acute myeloid leukaemia (AML) [4, 5]. This has led to the identification of several dysregulated signalling pathways in AML that may inform treatment.

It is important to note that racial differences in disease occurrence and treatment outcomes are well-known in medicine and have been under intense investigation for some time [6–8]. In the context of AML, ethnic and racial disparities have also been documented, influencing treatment outcomes and survival rates [9, 10]. Precision oncology and targeted therapies offer great promise for enhancing cancer outcomes; however, they are primarily based on biological shows a list of 197 enriched kinases in European AML patients with mechanisms and genetics that have not been thoroughly studied in minority races, including African populations [11]. Studies have revealed that African patients with AML have worse clinical outcomes and higher mortality rates compared to European patients [12, 13]. These findings serve as a call to action for better data collection and a deeper understanding of the genetic differences in AML among ethnic and racial groups. The impact of ethnic and racial disparities is not unique to AML, as findings in other cancers, such as those of the breast, have suggested a need for personalised management based on race and ethnicity [14, 15]. Thus, genetic factors may play a role in these disparities, as some mutations may be more prevalent in certain ethnic groups, thus affecting the treatment and survival outcomes of the afflicted patients.

In this study, we will leverage The Cancer Genome Atlas (TCGA) [16] clinical and transcriptomics datasets for AML obtained from cBioPortal [17] to conduct survival and functional enrichment analyses. We aim to investigate the disparities in survival outcomes and genetic profiles between European and African AML patients and to understand how these genetic variations influence drug responsiveness.

## Results

### Description of the datasets

We obtained a publicly available TCGA project [16] dataset from cBioPortal [17] for 200 AML patients, including clinical sample information with patient attributes and comprehensive transcriptomics data. Our study did not involve the recruitment of human subjects or animal studies. Comparing the mean ages of African and European patients, we observed a close distribution: 51 years for Africans and 55 years for Europeans (**Figure 1c**). In our analysis of gender distribution, we noted that 40% of African patients were female, whereas 55% of European patients were female.

**Figure 1:**
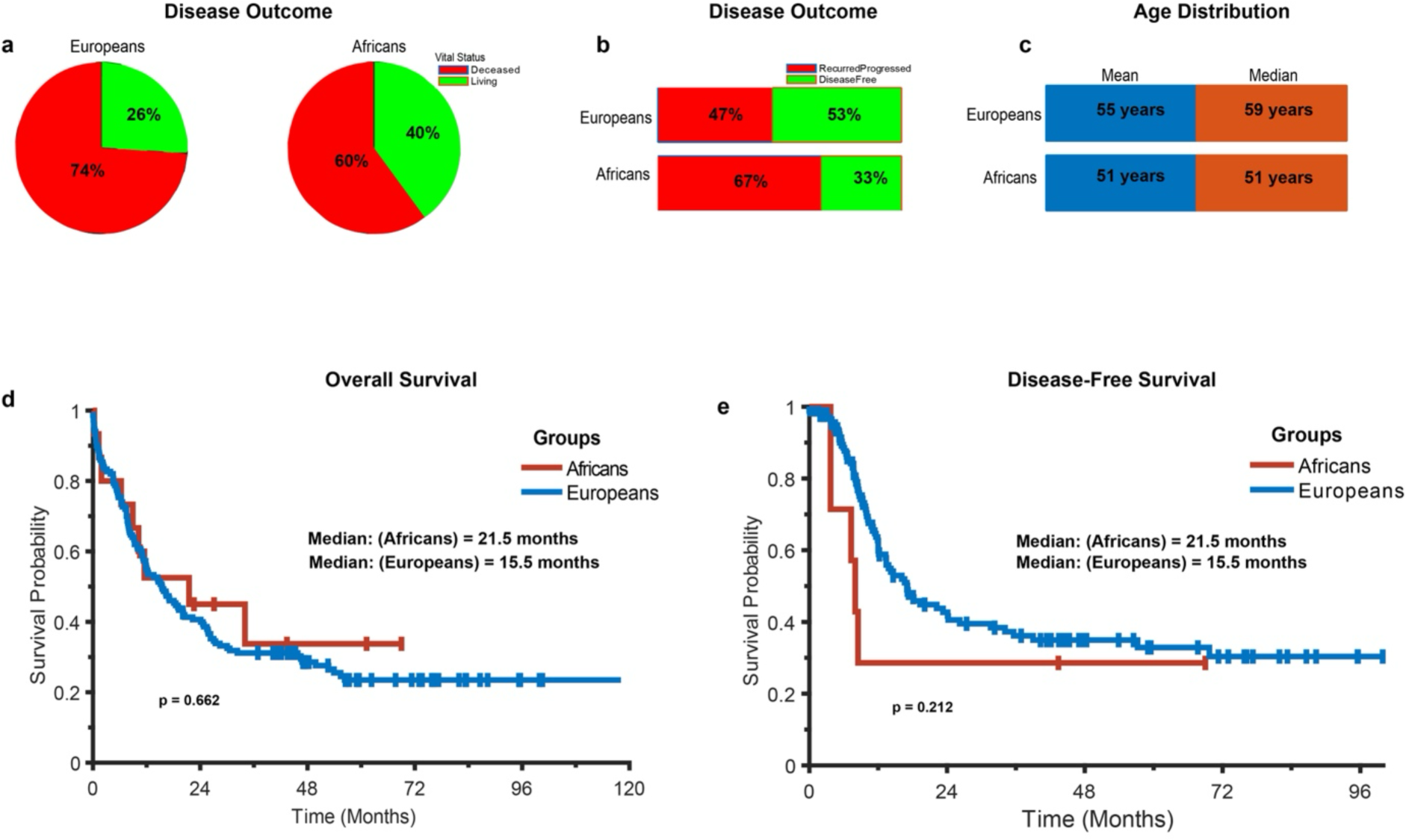
(a) Pie chart showing vital statistics after follow-up across the two races with AML; (b) Showing disease outcome after treatment; (c) Distribution of age across the two races with AML; (d) Kaplan-Meier curve for overall survival months for patients with AML across the two races; (e) Kaplan-Meier curve for Disease-Free Survival months for patients with AML across the two races.

### Clinical characteristics and survival outcomes of AML in Africans and Europeans

We aimed to evaluate any differences in survival outcomes between African and European individuals with AML. Thus, we compared the median disease-free survival (DFS) period and found that the median DFS duration was similar (Log-rank test; p = 0.212) for African patients (8 months) compared to European patients (17 months; **Fig 1e**). Additionally, we found comparable overall survival (OS) durations for African patients with AML (21.5 months) (p = 0.662) and European patients (15.5 months; **Fig 1d**).

We evaluated differences in other disease outcomes, including disease progression or recurrence at the end of follow-up in African and European individuals with AML. We found that 53% of European patients with AML were disease-free at the end of follow-up, while 47% experienced disease progression or recurrence, with only 26% surviving by the end of the follow-up period. In comparison, 33% of African patients with AML were disease-free at the end of follow-up, while 67% experienced disease progression or recurrence, with 40% surviving by the end of the follow-up period (**Figure 1 a and b**).

### Differentially expressed genes in AML between African and European patients

To comprehend the biological disparities between European and African individuals, we compared the differentially expressed genes in AML [18] between the two groups. We found 233 mRNA transcripts that were significantly upregulated in Europeans than in Africans, whereas 91 were significantly upregulated in Africans (**Figure 2a; Supplementary file 1**).

**Figure 2:**
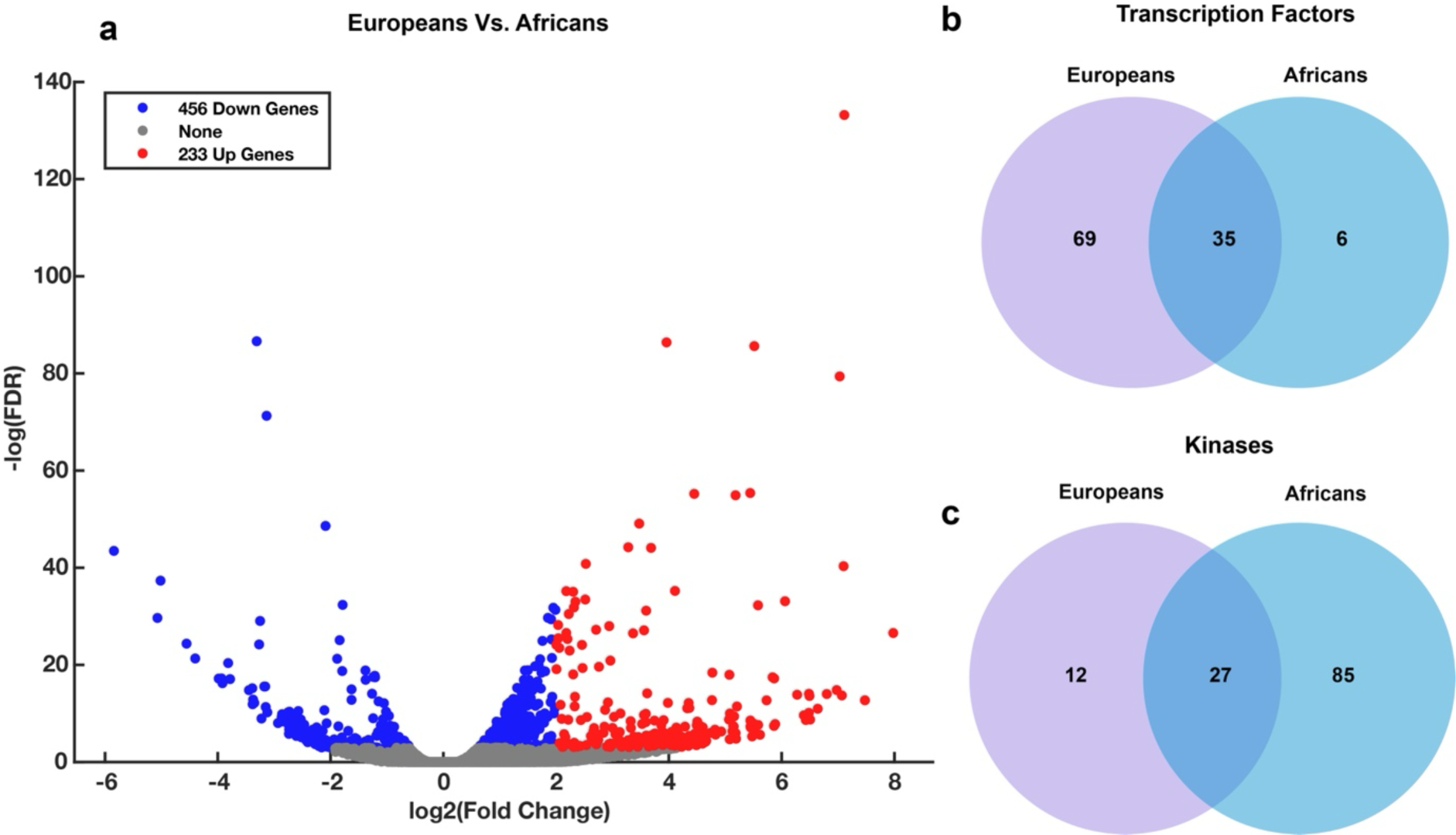
(a)Volcano plot showing 233 significantly upregulated and downregulated genes in Europeans with AML; (b) Venn diagram showing an overlap of 35 transcription factors across the two races, with 69 expressed significantly only in Europeans and 6 only in Africans; (c) Venn diagram showing an overlap of 27 kinases across the two races with 12 significantly expressed only in Europeans and 85 only in Africans

### Transcription factors and kinases distinguish AML in Africans and Europeans

Transcription factors are pivotal in the progression of cancer via their regulation and modulation of oncoprotein expression, including kinases. To understand gene expression and associated pathways in African and European individuals with AML, we utilised the expression-2-kinase software [19] to extract information regarding the transcription factors and kinases implicated in AML in these distinct racial groups.

Our analysis revealed considerable enrichment for several transcription factors, some shared between European and African individuals with AML, while others were specific to each racial group. Among the shared transcription factors were: SUZ12, SOX2 and POU3F2; Africans (p = 4.56 × 10^-12^, 6.36 × 10^-5^, and 7.28 × 10^-7^) and Europeans (p = 4.57 × 10^-29^, 2.61 × 10^-14^, and 4.65 × 10^-11^), respectively (**Figure 2b; Supplementary file 2**). Previous studies demonstrate the involvement of SUZ12 in the tumorigenesis of various cancers, including head and neck squamous cell carcinoma [20], endometrial and uterine cancer [21, 22], prostate cancer [23], and breast cancer [24]. Additionally, numerous studies indicate a correlation between high levels of SOX2 and poor prognosis in patients with various cancers, including breast, colon and rectum, oesophagus, ovaries, prostate, lung, and nasopharynx [25–27]. High levels of SOX2 are associated with regulating stem cell self-renewal [28] and the self-renewal and tumorigenicity of stem-like cancer cells [29, 30]. Furthermore, the transcription factor POU3F2 has also been documented to contribute to tumorigenesis in human gliomas in vivo [31].

Our findings indicate unique enrichment for TP53 (p = 2.64 × 10-09), RNF2 (p = 2.89 × 10-23), and EZH2 (p = 2.89 × 10-23) transcription factors in AML of European individuals. Conversely, significant enrichment was identified for SMAD4 (p = 0.001), EWS-FLI1 (p = 0.005), and CEBPB (p = 0.01), among others, only in AML of African individuals. RNF2 and EZH2 belong to the Polycomb-group (PcG) family, which has been demonstrated to regulate the expression and function of several oncogenes and tumour suppressor genes and has been linked to cancer patient survival [32]. Targeting the polycomb machinery, through pharmacological or genetic means, has been proposed as a promising approach for cancer therapy [32, 33].

Our kinase enrichment analysis showed substantial overlap in kinases in AML of African and European individuals and some kinases that were significantly enriched in AML of a specific racial group only. The top-ranked kinases commonly enriched in AML of both races included CNSK2A, AKT1, HIPK2, TAF1, MAPK1, and MAPK14, among others. (**Figure 2c; Supplementary file 3**)

Additionally, ULK2 (p = 0.013), CSNK2B (p = 0.021), and CAMK1 (p = 0.022) were only enriched in the AML of European individuals. Conversely, MAPK3 (p = 2.00 × 10-8), ATM (p = 1.93 × 10^-9^), CDK1 (p = 7.35 × 10^-5^), CDK4 (p = 4.20 × 10^-5^), CDK7 (p = 1.69 × 10^-6^), CDK8 (p = 0.007), CDK9 (p = 6.29 × 10^-6^), and CHEK1 (p = 2.62 × 10-^04^) were only significantly enriched in AML of Africans. Cyclin-dependent kinases (CDK1, 4, 7, 8, 9 and CHEK1) are crucial cell cycle regulators [34, 35]. Therefore, the significant enrichment of these kinases in AML of Africans may indicate substantial dysregulation of cell cycle processes. Hence, drugs targeting these kinases might eliminate AML cells and/or reduce their metastases [36, 37].

### Signalling pathways and molecular processes distinguish AML in Africans and Europeans

To determine functional terms enriched in AML for each race, we utilized Enrichr (https://maayanlab.cloud/Enrichr/) to query lists of up-regulated genes in AML among African and European individuals. Based on the WikiPathway and Bioplanet database terms, we found that AML in Europeans displayed significant enrichment for the Glial Cell-line Derived Neurotrophic Factors/ Rearranged during Transfection (GDNF/RET) signalling axis (p = 1.34×10^-4^), genes controlling nephrogenesis (p = 1.38 × 10^-4^), Gamma-Aminobutyric Acid (GABA) receptor activation (p = 8.26 × 10^-6^) and ligand-gated ion transport channels (p = 1.87 × 10^-4^), among others (**Fig 3a, b and d**). Previous studies have reported the upregulation of GABAergic signalling events in bone marrow lymphocytes in childhood acute lymphoblastic leukaemia [38]. In addition, GABAergic signal has also been found to play a proliferative effect on oral squamous cell carcinoma cell lines in association with mitogen-activated protein kinase signalling pathways [39]. Increased GDNF/RET expression or activity has been observed in various cancers, such as breast cancer [40, 41], pancreatic cancer [42], prostate cancer [43, 44], and myeloid malignancies [45, 46]. Moreover, high GDNF/RET levels have been linked to poor prognosis and survival, including tumour-related pain [42, 45, 47–49].

**Figure 3:**
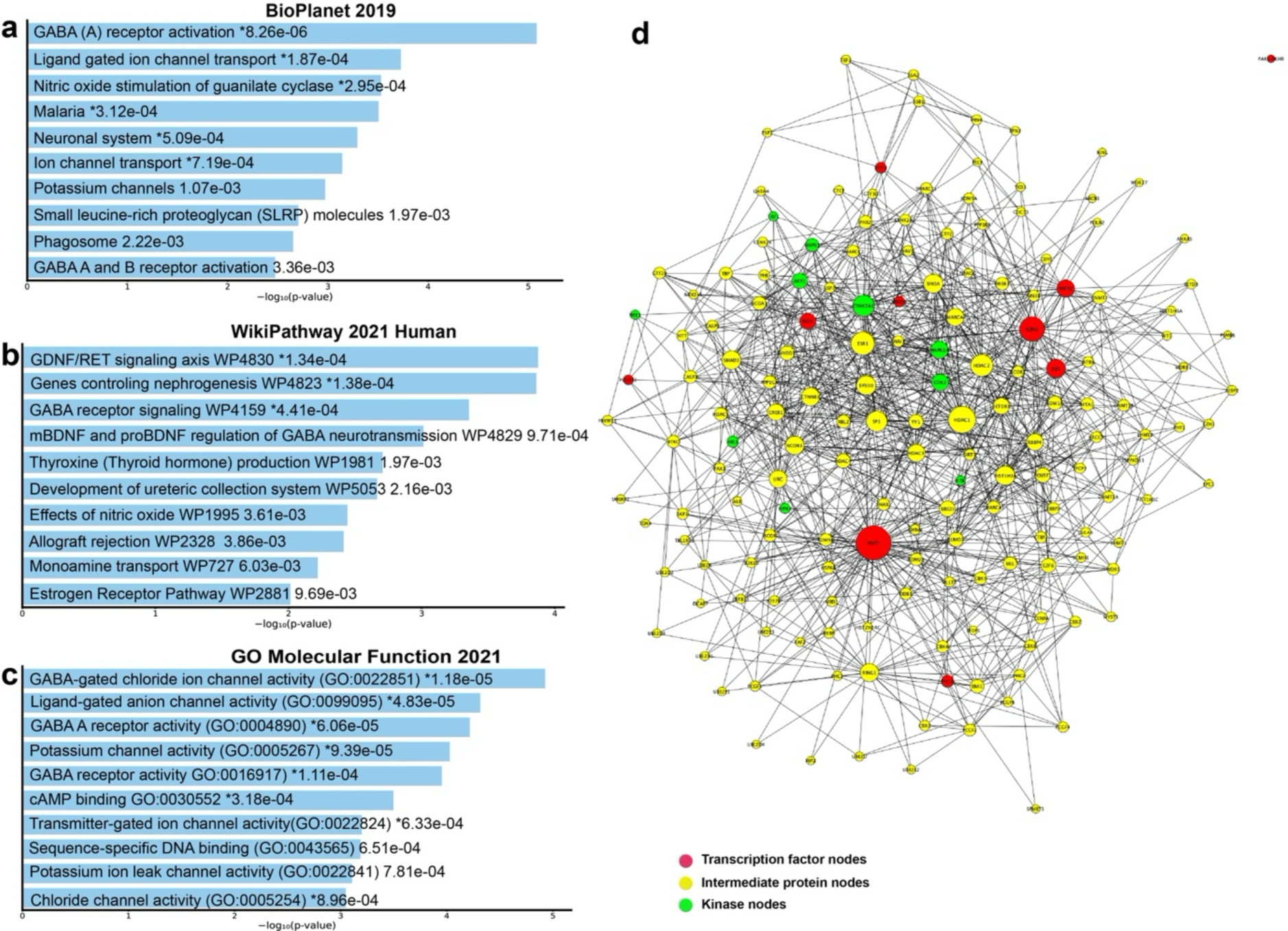
(a) Bar chart showing the top 10 enriched signalling pathway terms for Europeans with AML in the BioPlanet 2019 library along with their corresponding p-values; (b) Bar chart showing the top 10 enriched signalling pathway terms for Europeans with AML in the WikiPathway 2021 human library along with their corresponding p-values; (c) Bar chart showing the top 10 enriched Go molecular process terms for Europeans with AML in the GO Molecular Function 2021 library along with their corresponding p-values; (d) Intermediate protein-protein interaction subnetwork connecting transcription factors to the upstream regulatory proteins: the sub-network has 167 nodes with 964 edges. The network is enriched for co-regulators, kinases and transcription factors

In contrast, analysis of enriched functional terms in AML of African individuals, using the lists of up-regulated genes, revealed significant enrichment for several WikiPathway and Bioplanet terms, including signalling by G-Protein-Coupled Receptors (GPCR) (p = 4.64 × 10^-4^), Oncostatin M (p = 5.45 × 10^-4^), Codeine and Morphine metabolism (p = 2.07 × 10^-3)^ and Glucuronidation (p = 6.2 × 10^-3^) (**Fig 4a, b and d**). GPCR signalling has been implicated in several hallmarks of cancer, including proliferative signalling, evasion of growth suppressors, resistance to apoptosis, initiation of angiogenesis, and activation of invasion and metastasis [50]. Previous studies have reported a role for GPCR signalling in the leukemogenesis of AML [51] and its importance in B cell signalling, cell migration, proliferation, apoptosis, development, and function [52]. The signalling pathways in codeine and morphine metabolism form part of the opioid system, which has been implicated in the progression of various types of cancers and has been linked with tumour prognosis [53–57]. As such, opioid receptors have become targets of treatment for various cancers, including AML [58–60]. Oncostatin M (OSM) is considered a tumour-associated cytokine, with high concentrations detected in the serum and tumour sites of cancer patients, including in colon cancer [61], pancreatic cancer [62], myeloma [63], brain tumours [64], Chronic Lymphocytic Leukaemia [65]. OSM exerts its functions through the exploitation of multiple signalling pathways, including the JAK/STAT pathway, ERK1/ERK2, c-Jun N-terminal kinase (JNK), p38, protein kinase C delta (PKCδ), and the phosphatidylinositol-3-kinase (PI3K)/Akt pathways [66]. Therefore, inhibiting OSM generation and signalling may offer a therapeutic approach to AML treatment [63].

**Figure 4:**
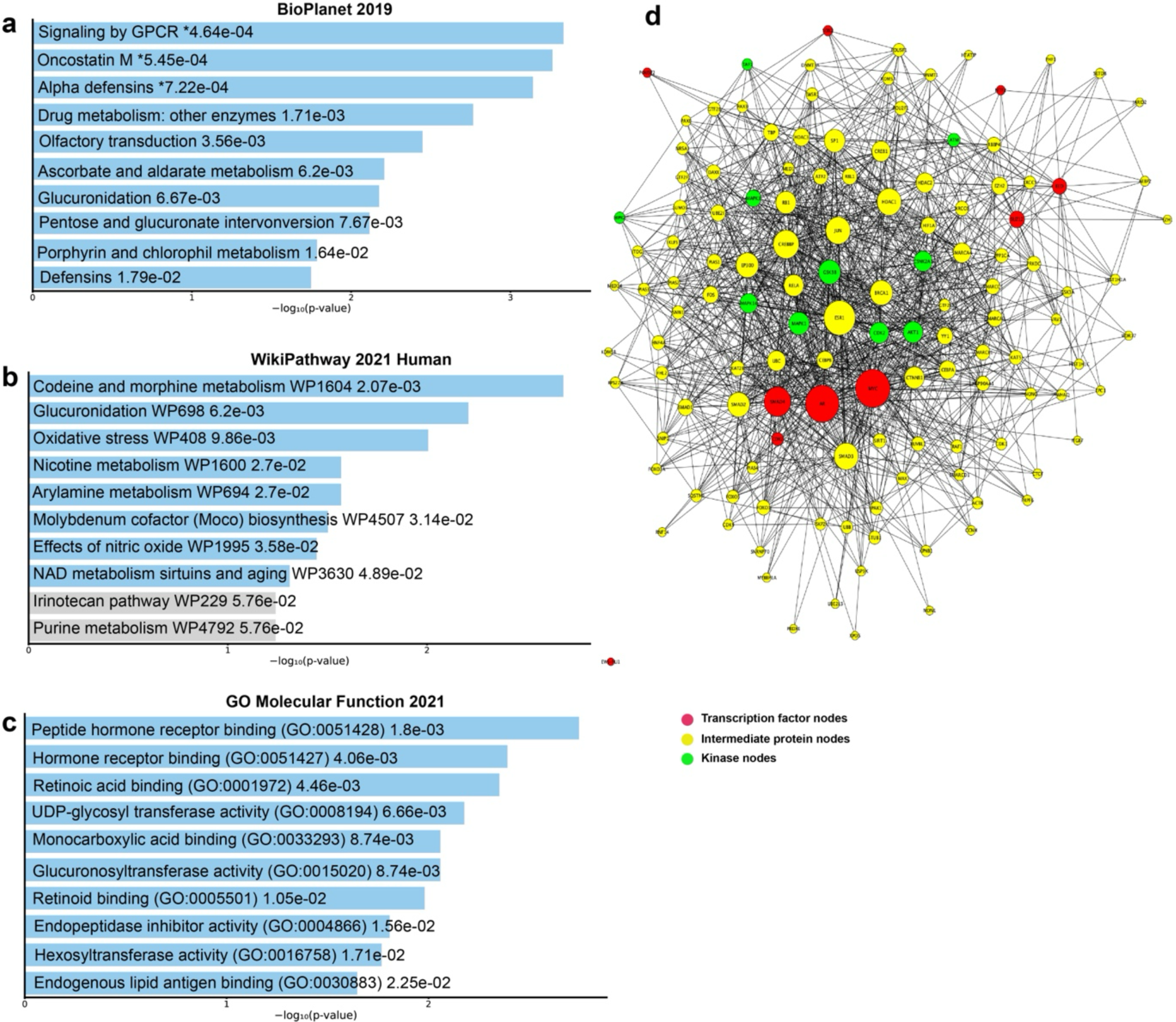
(a) Bar chart showing the top 10 enriched signalling pathway terms for Africans with AML in the BioPlanet 2019 library along with their corresponding p-values; (b) Bar chart showing the top 10 enriched signalling pathway terms for Africans with AML in the WikiPathway 2021 human library along with their corresponding p-values; (c) Bar chart showing the top 10 enriched Go molecular process terms for Africans with AML in the GO Molecular Function 2021 library along with their corresponding p-values; (d) Intermediate protein-protein interaction subnetwork connecting transcription factors to the upstream regulatory proteins: the sub-network has 130 nodes with 1179 edges. The network is enriched for co-regulators, kinases and transcription factors

To examine the biological processes involved in AML of Africans and Europeans, we performed a Gene Ontology (GO) [67, 68] analysis on the up-regulated genes. The molecular processes enriched in AML of Europeans were found to include GABA-gated chloride ion channel activity (p = 1.18 × 10^-5^), ligand-gated anion channel activity (4.83 x10^-5^), and GABA A receptor activity (p = 6.06 × 10^-5^) (**Fig 3c**). These results correspond with the WikiPathway and BioPlanet enrichments identified in the AML of Europeans.

Furthermore, our analysis revealed that the AML in African individuals was significantly enriched for molecular process terms in the Gene Ontology (GO) database associated with peptide hormone receptor binding (p = 1.8 × 10^-3^), hormone receptor binding (p = 4.06 × 10^-3^), and retinoic acid binding (p = 4.46 × 10^-3^), among others (**Figure 4c**). The processes of hormone receptor binding, primarily reported in breast cancer [69, 70], have been linked to drug resistance and patient outcomes [70]. Hormones acting through hormone receptors stimulate gene transcription and cell proliferation and are linked to the cell cycle machinery and cancer [71, 72]. Furthermore, it has been demonstrated that retinoic acid can cause cell cycle arrest and apoptosis in cancer cells [73]. Despite a mixed outcome across leukaemia subtypes, retinoic acid treatment has shown promise as a cancer therapy, with emerging biomarkers suggesting that certain subgroups may exhibit enhanced sensitivity [74]. Combination therapy, including retinoids is therefore a promising approach [75].

### mRNA Transcripts and Signalling Pathways Associated with Drug Response

We assessed whether the significantly differentially expressed mRNA transcripts in African and European AML patients correlated with variations in the chemosensitivity of AML cell lines to specific anticancer drugs. Through this approach, we intended to pinpoint ethnic-specific drug classes targeting specific, potentially enhancing AML treatment strategies for each group. Our findings demonstrated that cell lines with low and high mRNA transcript levels of the differentially expressed genes exhibited statistically significant differences in chemosensitivity to drugs targeting nine signalling pathways (Figures 5 and 6).

**Figure 5:**
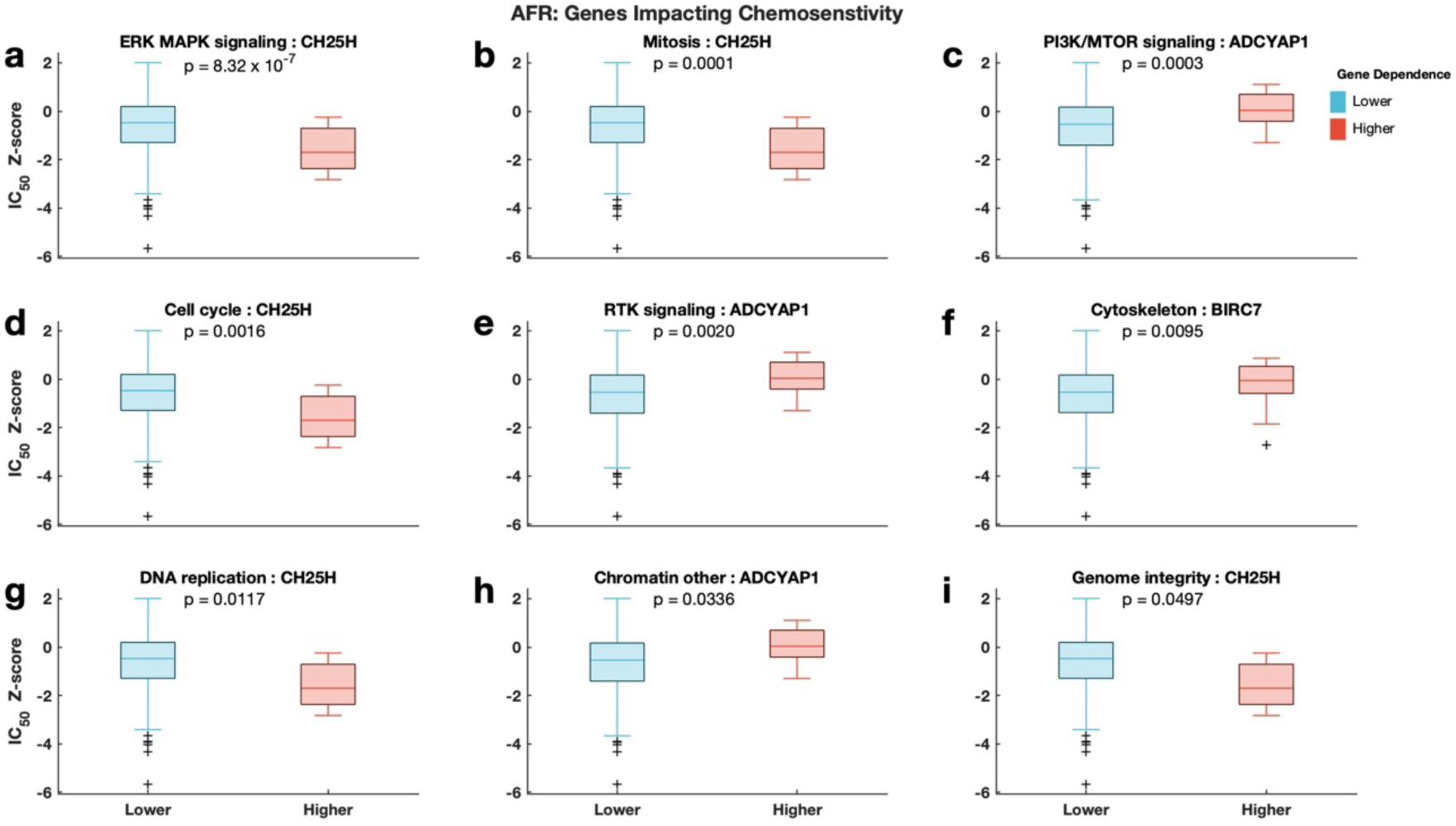
Showing mRNA transcript levels for genes involved in signalling pathways and how they relate to response to anti-cancer drugs in African AML patients. Cell lines expressing high levels of CH25H mRNA transcripts were more responsive to drugs targeting the (a) ERK/MAPK, (b) Mitosis, (d) cell cycle, (g) DNA replication and (i) genome integrity signalling pathway. However, those expressing low levels of ADAMTS19 gene mRNA transcripts were more sensitive to drugs that target the (c) PI3K/MTOR, (e) RTK and (h) cytoskeleton signalling.

**Figure 6:**
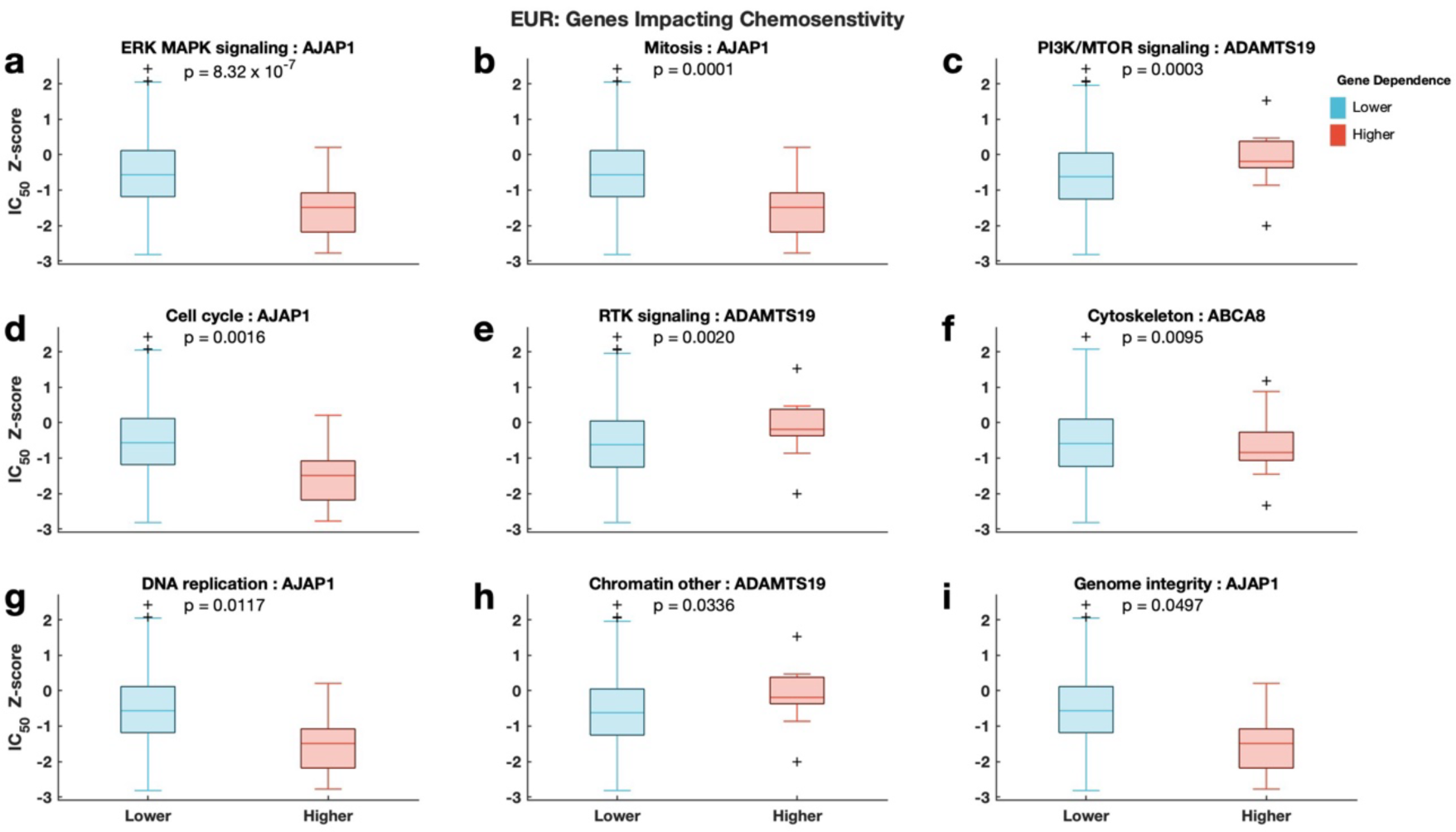
Showing mRNA transcript levels for genes involved in signalling pathways and how they relate to response to anti-cancer drugs in European AML patients. Cell lines expressing high levels of AJAP1 and ABCA8 mRNA transcripts were more responsive to drugs targeting the (a) ERK/MAPK, (b) Mitosis, (d) cell cycle, (f) cytoskeleton, (g) DNA replication and (i) genome integrity signalling pathway. However, those expressing low levels of ADCYAP1 and BIRC& genes mRNA transcripts were more sensitive to drugs that target the (c) PI3K/MTOR, (e) RTK and (h) chromatin signalling.

Of the 38 significantly upregulated mRNA transcripts in the African cohort, 97 showed variations in chemosensitivity of cell lines to drugs targeting specific signalling pathways: Cell cycle (25 transcripts), ERK MAPK signalling (25), Mitosis (25), PI3K/MTOR signalling (11) and RTK signalling pathway (11 transcripts). Conversely, from the 104 significantly upregulated mRNA transcripts in the European cohort, we found 287 associated with the chemosensitivity of cell lines to drugs targeting specific pathways. The breakdown is as follows: Cell cycle (79 transcripts), ERK MAPK signalling (79), Mitosis pathways (79), PI3K/MTOR signalling (25) and RTK signalling pathway (25 transcripts).

Specifically, for transcripts upregulated in African AML patients, we observed that cancer cells with overexpressed *CH25H* gene mRNA levels displayed heightened sensitivity to drugs targeting several signalling pathways: ERK/MAPK (p = 8.32 X 10-7), Mitosis (p = 0.0001), Cell cycle (p = 0.0016), DNA replication (p = 0.0117), and genome integrity (p = 0.0497). Furthermore, in cancer cell lines with increased mRNA expression of the *ADCYAP1* and *BIRC7* genes, we noticed decreased sensitivity to drugs targeting the PI3K/MTOR (p = 0.0003), RTK (p = 0.0020), cytoskeleton (p = 0.0095), and chromatin (p = 0.0336) signalling pathways (Figure 5). Increased expression of *BIRC7* has been extensively linked with disease progression, tumorigenesis, and undesirable clinical outcomes across many cancers, including Non-Hodgkin Lymphoma [76], extrahepatic cholangiocarcinoma [77] adult AML [78] and pancreatic ductal adenocarcinoma [79]. Moreover, the role of *BIRC7* is particularly pronounced in drug resistance, especially during bladder cancer treatments [80].

Similarly, *CH25H* has consistently emerged as a reliable prognostic marker in various cancer types. Lower expression levels of this gene are typically synonymous with heightened disease severity and a poor prognosis in lung adenocarcinoma [81] and pancreatic cancer [82].

Regarding *ADCYAP1*, reduced mRNA expression levels, often due to transcriptional silencing by hypermethylation, are associated with cervical carcinogenesis. This downregulated expression is prevalent in advanced stages of the disease and correlates with invasive carcinoma in cervical cancer [83]. In contrast, elevated *ADCYAP1* mRNA levels are linked to a shorter disease-free survival time in colorectal cancer patients [84].

For European AML patients, cancer cells overexpressing mRNA levels of the *AJAP1* and *ABCA8* genes, which were significantly upregulated, exhibited heightened sensitivity to drugs targeting the ERK/MAPK (p = 8.32 X 10^-7^), mitosis (p = 0.0001), cell cycle (p = 0.0016), cytoskeleton (p = 0.0095), DNA replication (p = 0.0117), and genome integrity (p = 0.0497) signalling pathways. In contrast, cells under expressing mRNA levels of the *ADAMTS19* gene displayed increased sensitivity to drugs that target the PI3K/MTOR (p = 0.0003), RTK (p = 0.020), and chromatin (p = 0.0336) signalling pathways (Figure 6) (**Figure 6**). Increased expression of *AJAP1* correlates with a favourable prognosis in breast cancer [85], and salivary adenoid cystic carcinoma [86] and augments the sensitivity of glioblastoma cells to temozolomide by promoting apoptosis [87]. *ABCA8*, part of the ATP Binding Cassette Subfamily A, when overexpressed, has ties to a favourable prognosis and curtails the proliferation and metastasis of hepatocellular carcinoma and breast cancer cells [88, 89]. Yet, the overexpression of *ABCA8* is linked to enhanced chemoresistance in several cancers, including pancreatic cancer and AML [90, 91]. Furthermore, the increased expression of *ADAMTS19* is associated with better overall survival and suppresses cell migration and invasion in human gastric cancer [92].

## Methods

We obtained a dataset of 200 acute myeloid leukaemia patients from the TCGA project [16] via cBioPortal (http://www.cbioportal.org) [17]. Our analysis was based on the clinical and mRNA gene expression data of 165 African and European AML patients, which was thoroughly de-identified to protect the anonymity of the patients. In the TCGA, the terminology utilised for races was ‘Black’ and ‘White’. However, for our research, we have chosen to use the terms ‘African’ for Blacks and ‘European’ for Whites.

### Survival Analysis

We employed the Kaplan-Meier method [93] to assess the differences in overall survival and disease-free survival between African and European patients, using the MatSurv software [94] in MATLAB.

### Identification of the differentially expressed genes

We used mRNA expression data to identify the differentially expressed genes between Europeans and Africans afflicted with AML. We conducted statistical analysis on the mRNA transcripts of both groups, utilizing the Negative Binomial Model [18]. The p-values obtained from the analysis were adjusted with the FDR method [95]. mRNA expression was considered statistically significantly differentially expressed in AML between Africans and Europeans when it exhibited an adjusted p-value of less than 0.05 and a fold-change greater than 4.

### Functional enrichment analyses

We used the list of significantly up-regulated genes in both Europeans and Africans separately to query Enrichr [96–98] for enriched signalling pathways and functional terms (BioPlanet [99], WikiPathway [100], and GO molecular function [67, 68, 101]) in AML of either Europeans or Africans. After identifying the functional enrichments, we used the yEd [102] network visualisation software to visualise the connectivity of components in each enriched term and pathway.

### Transcription factor and kinase enrichment analysis

We employed the X2K software [19] to infer transcription factors and kinases from the lists of differentially expressed genes. We ran the lists of significantly upregulated genes in Africans and Europeans independently through the X2K software and extracted statistically significant transcription factors and kinases from the .csv file outputs.

### Determination of Genes and Signalling Pathways Associated with Drug Response

To identify genes linked with notable variations in the dose-response profiles of cancer cell lines, we turned to the lists of genes differentially expressed for each ethnicity from our preceding analysis. We extracted mRNA expression details for these genes from the CCLE expression data. We then categorised these expression levels into ‘low’ or ‘high’ expression brackets. Following this, we extracted the drug-response profile for these cell lines from the GDSC, which covered 304 anticancer drugs targeting 23 pathways. We then employed the Welch Test to determine whether these mRNA transcripts were associated with significant variations in the dose-response patterns of AML cell lines.

### Statistical analysis and reproducibility

We employed MATLAB version 2022b for conducting all the analyses presented in this study. Descriptive statistics were utilized to characterize the age and sex distribution of participants. Statistical significance was established when p-values were less than 0.05.

## Discussion

In this study, we conducted a comprehensive analysis to investigate the differential gene expression between African and European patients with AML. Our results demonstrate the presence of distinct differences in the genetic landscape of AML patients of the two races, indicating potential disparities in the pathogenesis and therapy response of AML between the two groups. These findings provide valuable insights to inform the development of more personalised treatment strategies for AML in African and European patients.

We demonstrated that survival outcomes (disease-free and overall survival) were comparable between the two races, but the median disease-free survival was lower for African patients than European patients. This aligns with many studies that have reported poor survival outcomes for African patients compared to European patients with AML [103, 104]. The minority races tend to be underrepresented in most of the studies compared to the non-Hispanic European participants relative to cancer incidence [11, 105–107]. Therefore, greater efforts are needed to diversify cancer drug trials to improve equity in access to new treatments and to ensure that safety and efficacy findings from early drug trials are generalizable across races [105, 108, 109].

At the end of follow-up, only 33% of African patients were disease-free, compared to 53% of European patients. This disparity may be attributed to the enrichment of the OSM pathway in African patients. The OSM pathway operates through various signalling pathways, including tumour cell survival pathways like ERK1/ERK2 and the PI3K/Akt pathways [66]. Therefore, the higher survival rate of cancer cells in Africans may explain why fewer African patients achieve disease-free status compared to European patients at the end of the follow-up period. Nevertheless, suppression of disease via mechanisms involving SMAD4 [110] which has been linked to multiple favourable prognostic factors and is associated with improved overall survival [111] might explain the higher survival of African AML patients by the end of follow up compared to their European counterparts.

AML development is a multi-cause, multi-step, and multi-pathway process and constitutes a heterogeneous group of diseases [112]. Our study has uncovered disparities in the signalling pathways and Gene Ontology molecular process terms involved in the pathogenesis of AML between European and African patients. Recent advancements in next-generation sequencing and multi-omics analysis have shed light on the intricacies of cellular signalling pathways and their potential for targeted therapy if fully understood [113, 114]. In light of these findings, we propose that precision medicine be employed in treating AML in different racial populations.

While some similarities were present in the expression of transcription factors and kinases, significant differences were observed between the two racial groups. The most striking difference was the enrichment of Cyclin-Dependent Kinases (CDKs) in AML of African patients. CDKs are critical in cancer, have prognostic value, and are treatment targets in many cancers. Previous studies have shown that high mRNA levels of certain CDKs are linked to poor prognosis and low overall survival and relapse-free survival in human breast cancer [115], gastric cancer [116], lung cancer [117], acute lymphoblastic leukaemia [118] and many other cancers. In recent years, CDKs have also become targets of inhibition in cancer therapeutics [119–123].

Molecular changes in cancer genes and their related signalling pathways guide the development of precision medicine treatments for cancer [124]. In this study, we analysed the link between mRNA transcript differences in African and European AML patients and AML cell line chemosensitivity. Among the African cohort’s upregulated mRNA transcripts, notably, the *CH25H* gene exhibited heightened sensitivity to drugs across several pathways, while *ADCYAP1* and *BIRC7* showed reduced sensitivity in specific pathways (Figure 5). Conversely, in the European cohort, *AJAP1* and *ABCA8* overexpression influenced sensitivity in pathways like ERK/MAPK, whereas *ADAMTS19* under expression affected others (Figure 6). Here, our findings underscore the importance of ethnic-specific molecular understanding for enhancing AML treatments.

This study sheds light on the disparities in molecular signatures and gene expression in response to chemotherapy between Europeans and Africans with AML. Our findings suggest that African patients have an enrichment of gene ontology molecular processes and kinases related to cell cycle regulation and differences in signalling pathways. These results indicate the potential for race-specific precision treatment in AML patients.

### Ethics approval

The protocol for this study was approved by the University of Zambia Biomedical Research Ethics Committee IRB00001131 of IORG0000774, approval number 3062-2022. The publicly available datasets were collected by the cBioPortal and TCGA projects and made available through their respective databases. All methods used were performed within the stipulated guidelines provided by the cBioPortal and TCGA projects.

## Supporting information

Supplementary File Description

## Conflicts of interest

Authors declare no conflicts of interest

## Author Contribution

**Conceptualisation:** Panji Nkhoma, Sinkala Musalula, Kevin Dzobo, Geoffrey Kwenda, Sody Munsaka

**Data curation**: Panji Nkhoma, Sinkala Musalula, Kevin Dzobo, Doris Kafita, Sody Munsaka

**Formal Analysis**: Panji Nkhoma, Sinkala Musalula, Kevin Dzobo, Doris Kafita, Geofrey Kwenda, Sody Munsaka

**Methodology**: Panji Nkhoma, Sinkala Musalula, Kevin Dzobo, Doris Kafita

**Visualisation**: Panji Nkhoma, Sinkala Musalula, Geoffrey Kwenda

**Writing – original draft**: Panji Nkhoma, Sinkala Musalula, Geoffrey Kwenda

**Writing – review & editing**: Panji Nkhoma, Sinkala Musalula, Kevin Dzobo, Doris Kafita, Geoffrey Kwenda, Sody Munsaka

## Notes

### Competing Interest Statement

The authors have declared no competing interest.

