## Supplementary File Description for "Racial Disparities in the Genetic Landscape of Acute Myeloid Leukaemia from The Cancer Genome Atlas: Insights from a Bioinformatics Analysis"

**Supplementary File 1:**

Supplementary file 1 is an xlsx file with three tabs showing differentially expressed genes in Europeans and Africans with Acute Myeloid Leukaemia (AML) generated from results of the Negative Binomial Test. ***EUR Vs AFR:*** shows a list of all the 20518 genes from the TCGA with mean mRNA expression values, fold change and p values**, *Up Genes EUR:*** shows a list of 233 upregulated genes in European AML patients with mean mRNA expression values, fold change and p values, ***Up Genes AFR:*** shows a list of 91 upregulated genes in African AML patients with mean mRNA expression values, fold change and p values.

**Supplementary File 2**

Supplementary file 2 is an xlsx file with two tabs showing enriched transcription factors enriched in European and African patients with AML. ***EUR:*** shows a list of 223 enriched transcription factors in European AML patients among the most enriched being JARID2, SUZ12, RNF2, EZH2 and MTF2, ***AFR:*** shows a list of 177 enriched transcription factors in African AML patients among the most enriched being SUZ12, POU3F2, MTF2, MYC and CDX2.

**Supplementary File 3**

Supplementary file 3 is an xlsx file with two tabs showing enriched kinases in European and African AML patients. ***EUR:*** shows a list of 197 enriched kinases in European AML patients with MAPK14, CSNK2A1, AKT1, HIPK2 and WEE1 being among the most enriched, ***AFR:*** shows a list of 241 enriched kinases in African AML patients with MAPK1, AKT1, MAPK3, MAPK14 and GSK3B among the most enriched
